# *Aedes aegypti* from a Population Co-occurring with *Aedes albopictus* Show Species-Selective Premating and Postmating Resistance to Reproductive Interference

**DOI:** 10.64898/2026.07.16.738986

**Authors:** Lauren E. Subramaniam, Olivia M. Martinez, Thomas M. Gabel, Laura B. Duvall

## Abstract

**Background:** A female *Aedes aegypti* mosquito typically mates only once, receiving all of the reproductive materials required for her lifetime from a single male. During mating, males transfer chemical signals that permanently suppress female receptivity to subsequent mating attempts. Successful mating can be disrupted by satyrization, a form of interspecific reproductive interference in which *Aedes albopictus* males inseminate *Aedes aegypti* females without producing viable offspring, effectively rendering them sterile. *Aedes aegypti* populations from areas where both species co-occur can develop resistance to satyrization, which may arise through premating mechanisms including altered mating interactions and/or postmating mechanisms including reduced female sensitivity to male-derived chemical compounds.

**Methodology/Principal Findings:** We compared two *Aedes aegypti* strains derived from field collections in Lee County, Florida: one from a site where *Aedes albopictus* is established (Fort Myers) and one from a site with no known history of *Aedes albopictus* co-occurrence (Upper Captiva Island). Using laboratory mating assays, fluorescent dye transfer experiments, and male reproductive homogenate injections, we show that females from the co-occurring population exhibit reduced susceptibility to satyrization. This resistance is associated with two distinct mechanisms: reduced physical interactions with heterospecific males and reduced physiological sensitivity to *Aedes albopictus* homogenate following injection. Notably, co-occurring females retain sensitivity to signals from conspecific males, indicating that resistance mechanisms are species-selective rather than a generalized reduction in mating responsiveness.

**Conclusions:** *Aedes aegypti* females from a population co-occurring with *Aedes albopictus* show reduced susceptibility to satyrization under laboratory conditions, mediated by both premating and postmating mechanisms. The species selectivity of these responses suggests that co-occurring populations can discriminate between conspecific and heterospecific signals. Understanding how this discrimination is achieved and how it varies across populations will be important for interpreting interspecific mating dynamics and for evaluating vector control strategies that depend on mating interference, including sterile male release programs.

## INTRODUCTION

*Aedes aegypti*, the yellow fever mosquito, and *Aedes albopictus*, the Asian tiger mosquito, are aggressive vectors that are responsible for the spread of disease-causing pathogens including dengue, chikungunya, yellow fever, and Zika virus [1]. Both species have spread globally through close association with human hosts: *Aedes aegypti* expanded from sub-Saharan Africa beginning in the 15th century, and *Aedes albopictus* spread from Asia in the mid-1980s [2,3]. Their ranges continue to expand, and climate change is predicted to accelerate this process [4,5]. By the year 2100, most people in the United States are projected to live in areas where both species co-occur, exposing new populations to vector-borne disease risk [1].

Where their ranges overlap outside of their native habitats, *Aedes albopictus* frequently displaces established *Aedes aegypti* populations [6–8]. Such displacements have been documented across Florida since the 1980s and in Bermuda, where the arrival of *Aedes albopictus* in the year 2000 was followed by a rapid decline in *Aedes aegypti* abundance [9,10]. Although *Aedes albopictus* larvae often outperform *Aedes aegypti* in low-nutrient environments, the speed of these displacements suggests that resource-based competition alone cannot explain these patterns [11–13]. Instead, satyrization, a form of interspecific mating interference, has been proposed as a key driver [6–8,10,13,14].

During mating, male *Aedes* mosquitoes transfer Seminal Fluid Proteins (SFPs), contained in Male Accessory Gland (MAG) fluids, to females. These proteins induce female refractoriness to subsequent mating, thereby enforcing the male’s paternity of all offspring [15,16]. Although females generally mate only once, males can inseminate multiple females. Satyrization occurs when interspecific mating imposes a fitness cost, in this context when *Aedes albopictus* males mate with *Aedes aegypti* females and transfer paternity-enforcing signals [17,18]. Because *Aedes albopictus* SFPs retain paternity-enforcing activity in *Aedes aegypti* females [19], satyrized females reject subsequent conspecific mates but are inseminated with incompatible sperm and produce inviable eggs, effectively sterilizing them [20]. This interaction is asymmetric between these two species and MAG fluids from *Aedes aegypti* males do not enforce paternity in *Aedes albopictus* females [10], providing support for the idea that satyrization contributes to the displacement of *Aedes aegypti* by invading *Aedes albopictus*.

Despite early displacement, *Aedes aegypti* populations have resurged in parts of Florida, suggesting the development of satyrization resistance. This resistance can manifest as reduced mating acceptance and/or increased ability to remate after interspecific mating [7,10,13]. Mating reluctance has been observed in field-collected populations of *Aedes aegypti* that co-occur with *Aedes albopictus* [13] and females inseminated by both species have been detected in the field, indicating that postmating recovery from satyrization occurs in nature [8]. These resistance traits are plastic; laboratory studies show that interspecific mating rates decline after only three generations of co-rearing, but mating avoidance can be quickly lost when resistant populations are reared without *Aedes albopictus* [14,21].

We examine two mechanisms by which *Aedes aegypti* females may resist satyrization: premating resistance, in which females make fewer physical contacts and are inseminated less often by *Aedes albopictus* males, and postmating resistance, in which females retain receptivity to conspecific males following exposure to *Aedes albopictus* SFPs. We hypothesize that *Aedes aegypti* females from populations that co-occur with *Aedes albopictus* experience selective pressure favoring these resistance mechanisms, whereas females from populations with no prior exposure do not.

Consistent with this hypothesis, females from a co-occurring population in Fort Myers, FL show fewer physical interactions with *Aedes albopictus* males and reduced sensitivity to *Aedes albopictus* MAG homogenate injection compared to females from a naive population on Upper Captiva Island, FL. These data indicate that *Aedes aegypti* females from the Fort Myers population possess both premating and postmating resistance mechanisms that reduce susceptibility to satyrization relative to females from the naive population.

## METHODS

### Field Collection

Field-derived populations of *Aedes aegypti* were established from eggs collected in oviposition traps placed in Lee County, FL between July 8 and July 17, 2024. Trapping sites were selected for their suitability for container-breeding *Aedes* mosquitoes, determined by proximity to human habitation and presence of artificial containers where rainwater could collect. Sites were selected in Fort Myers, where both *Aedes aegypti* and *Aedes albopictus* have longstanding co-occurrence, and Upper Captiva Island, where *Aedes albopictus* has never been detected in records sourced from Lee County Vector Control. Sites were not treated with larvicides or adulticides when the traps were in use. Eggs used to generate the Fort Myers (co-occurring) strain were collected from public land in the Ortiz neighborhood, Fort Myers Cemetery, and one private residence near downtown Fort Myers. Eggs used to generate the Upper Captiva (naive) strain were collected from two construction sites and four private residences on Upper Captiva Island. GPS coordinates for all collection sites are provided in **Table 1**.

**Table 1:** Summary of collection sites used to generate field-collected strains from Fort Myers and Upper Captiva.

| Population | Site Name | GPS Coordinates | *Ae. aegypti*<br>present | *Ae. albopictus*<br>present |
| --- | --- | --- | --- | --- |
| Fort Myers | Ortiz 1 | 26.645722, -81.810750 | x | x |
| Fort Myers | Ortiz 2 | 26.646278, -81.807750 | x | x |
| Fort Myers | Ortiz 3 | 26.652694, -81.809361 | x | x |
| Fort Myers | Ortiz 4 | 26.656056, -81.811944 | x | x |
| Fort Myers | Ortiz 5 | 26.652778, -81.807694 | x | x |
| Fort Myers | Cemetery | 26.649111, -81.848278 | x | x |
| Fort Myers | Private Residence | 26.658250, -81.847500 | x | x |
| Upper Captiva | Construction 1 | 26.597417, -82.219250 | x |  |
| Upper Captiva | Private Residence 1 | 26.595500, -82.220000 | x |  |
| Upper Captiva | Private Residence 2 | 26.595694, -82.221056 | x |  |
| Upper Captiva | Private Residence 3 | 26.595639, -82.221917 | x |  |
| Upper Captiva | Construction 2 | 26.602667, -82.218694 | x |  |
| Upper Captiva | Private Residence 4 | 26.603694, -82.218056 | x |  |

Each oviposition trap consisted of a 475 mL black plastic cup filled approximately halfway with deionized (DI) water. A paper towel was placed in the cup so that it was partially submerged and soaked, with one end overhanging the rim of the cup and secured in place with a heavy-duty rubber band. A hole was drilled in each cup approximately 2 cm below the rim to allow for overflow, so that eggs would not be lost in the event of rain. At each trapping site, traps were placed in locations that were shaded to prevent evaporation but not wholly covered to remain accessible to gravid females. Traps were left in the field for 3 days, after which papers with eggs were collected and traps were reset for another 3 days. Papers were hung to air-dry for several days (until completely dry), then packaged in resealable plastic bags and transported to the laboratory.

### Establishment of Field-Derived Strains

Eggs were hatched by placing papers from the same site in 1 L of hatch broth (deoxygenated DI water with finely-ground Tetramin tropical fish food) and larvae were reared until pupation. Pupae were collected by hand and transferred into a small cup of DI water in an insect rearing cage (BugDorm) cage until eclosion. Cages with pupae were examined twice per day for newly-eclosed adults, which were identified to species and separated into an *Aedes aegypti* or *Aedes albopictus* cage. No other species were found in our traps. Cohabitation for the Fort Myers sites was confirmed by the presence of both *Aedes aegypti* and *Aedes albopictus* eggs at all trapping sites, while no *Aedes albopictus* were found in any traps from Upper Captiva (see **Table 1**).

### Rearing

All mosquitoes were maintained and reared at 28°C, 70–80% relative humidity (RH) with a 12 h light: 12 h dark schedule [22]. Three strains were used in this study: one *Aedes albopictus* laboratory reference strain (Foshan; FOS) [23,24] and two field-derived *Aedes aegypti* strains (Fort Myers and Upper Captiva) (**Figure 1A**). The Fort Myers strain (FtM) was collected from a population that co-occurs with *Aedes albopictus*, and is therefore expected to have experienced selective pressure for satyrization resistance. The Upper Captiva strain (UpC) was collected from a population with no known history of co-occurrence with *Aedes albopictus*, and is considered naive with respect to interspecific mating pressure. Field-derived *Aedes aegypti* used in behavioral experiments were from laboratory generations F3-F6. Eggs were hatched in Hatch Broth (deoxygenated DI water with finely-ground Tetramin tropical fish food) and raised on a diet of Tetramin tropical fish food tablets or flakes (Tetra, Melle, Germany). Larval density was adjusted to approximately 250 individuals per rearing pan (17 × 16 × 10 cm). Animals were separated by sex at the pupal stage and housed separately as adults to ensure all animals were unmated for behavioral assays. Adults were provided 10% sucrose *ad libitum*. Adults were 3-10 days posteclosion at the time of experiments to ensure sexual maturity. To confirm the accuracy of sex separation and ensure that all females were unmated at the start of behavioral experiments, each experimental replicate was run concurrently with a cohort of females maintained in a “female only” cage with no male exposure. Any replicate in which insemination was detected in this control cohort was excluded from analysis. For colony maintenance, adult females were blood-fed using an artificial membrane feeder (Hemotek, UK) supplied with defibrinated sheep blood (Quad Five) supplemented with 2 mM ATP (Sigma-Aldrich). A section of nylon fabric previously worn by an experimenter was stretched over the feeding membrane to provide human olfactory cues and improve attraction to the feeder.

**Figure 1:**
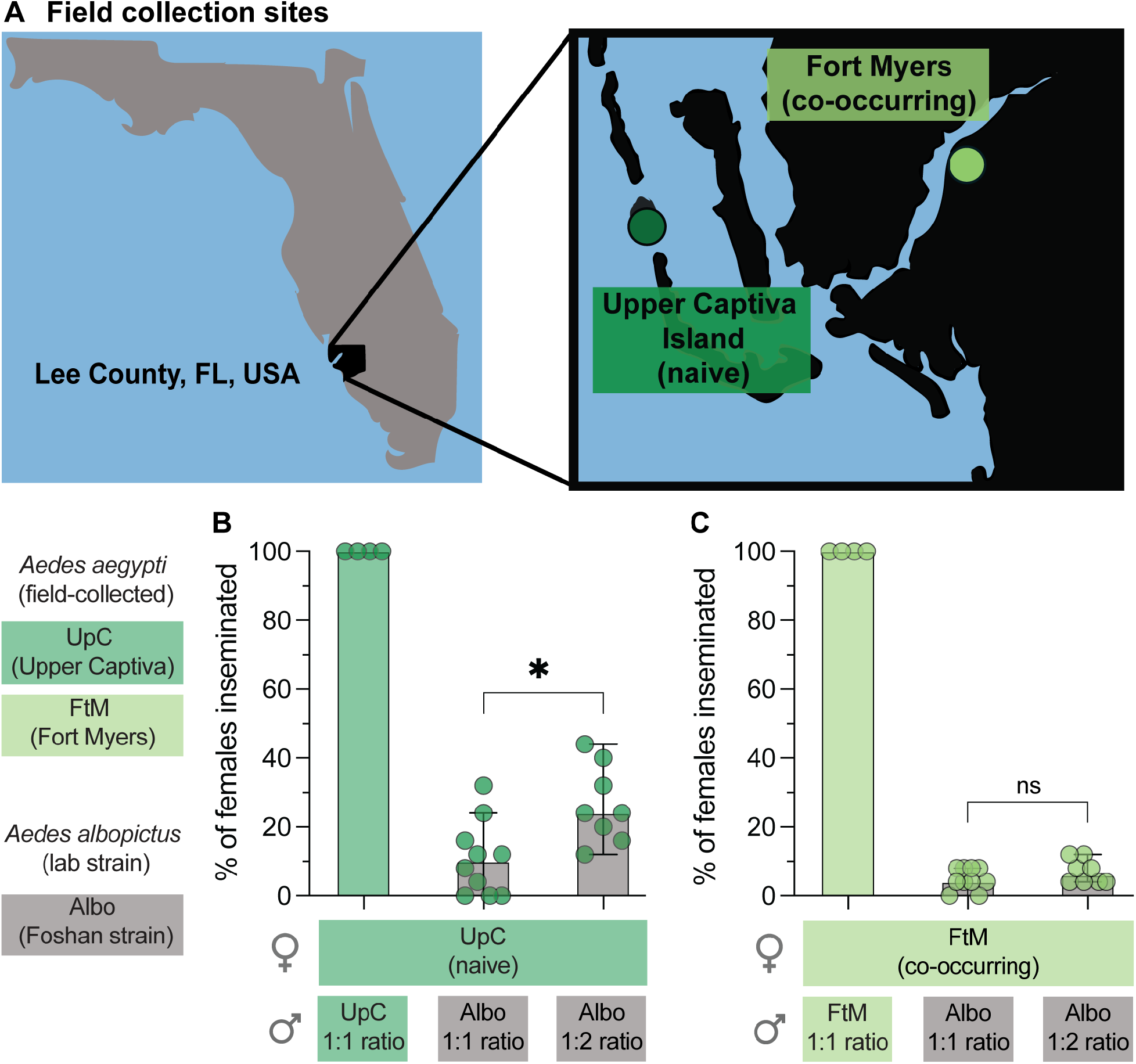
Field collection sites and strain susceptibility to interspecific insemination. (A) Map of *Aedes aegypti* field collection sites in Lee County, FL, USA. The Upper Captiva Island site (UpC) has no known history of *Aedes albopictus* co-occurrence; the Fort Myers site (FtM) is inhabited by both species. (B) Percentage of *Aedes aegypti* females from the naive UpC strain inseminated by conspecific (*Aedes aegypti*) or heterospecific (*Aedes albopictus*) males at 1:1 and 1:2 (female:male) ratios. (C) Percentage of *Aedes aegypti* females from the co-occurring FtM strain inseminated by conspecific or heterospecific males at 1:1 and 1:2 ratios. For all panels, n = 4-8 replicates with 23-25 females per replicate, drawn from at least 2 hatches. Asterisks indicate statistically significant differences (p < 0.05, Mann-Whitney U test; ns, not significant). Additional statistical details provided in **Data File S1**.

### Group Insemination Assay

To assess insemination rates, unmated females and males aged 3-10 days posteclosion were cohoused in large rearing cages (BugDorm) for 7 days at either a 1:1 or 1:2 (female:male) ratio (25 females paired with 25 or 50 males, respectively). In intraspecific trials, females were housed with *Aedes aegypti* males of the same strain; in interspecific trials, females were housed with *Aedes albopictus* males from the Foshan laboratory strain. The 1:2 female-to-male density was tested only in interspecific trials, as intraspecific trials at the 1:1 ratio yielded 100% insemination. Replicates were drawn from at least two different egg hatches in each experiment. Survivorship was recorded daily; dead males were replaced to maintain the target sex ratio, and males were removed proportionally if females died. Any replicate in which female mortality exceeded 20% was excluded from analysis. Following 7 days of cohabitation, all females were dissected and spermathecae were examined to determine insemination status.

### Dye Transfer Assay

Male mosquitoes were cold anesthetized at 4°C for 10 minutes, aspirated into a plastic cup, and placed on ice before dye application. A fluorescent oil-based dye (ACDelco 1,148,963 GM Original Equipment 10-5045 Multi-Purpose Fluorescent Leak Detection Dye) was applied to the terminal two segments of each male, as previously described [25]. Males were allowed to recover for one hour at 25-28°C. Individual males were aspirated into small cages (21.6 cm diameter x 18.1 cm height) each containing 10 unmated *Aedes aegypti* females. After 22 hours of co-housing, males were removed and females were cold anesthetized at 4°C for 10 minutes. Dye transfer to females was assessed under a Leica MZ10 F fluorescence stereomicroscope using the Cy3 channel. Insemination status was confirmed independently by spermathecal dissection.

### Male Abdominal Tip (MAT) Homogenate Preparation

Male mosquitoes were cold anesthetized at 4°C for 10 minutes, aspirated into a plastic cup, and placed on ice before MAT collection. Under a dissecting stereomicroscope, the terminal two abdominal segments of each male (containing the male accessory glands) were collected using Vannas-Tübingen spring scissors and transferred to a 1.5 mL microcentrifuge tube containing 10 µL of 1x PBS per 50 MATs and stored at -80°C until homogenization.

Before homogenization, each tube of MATs was diluted with 1x PBS to achieve a concentration of 1 µL 1X PBS per MAT. MATs were homogenized in the microcentrifuge tube using a motorized pestle grinder (Kimble Pellet Pestle Cordless Motor 749540-0000) for 1 minute and briefly centrifuged to pellet physical debris. The supernatant containing MAT homogenate was then transferred to a 0.45 µM centrifugal filter (Durapore PVDF 0.45 μM UFC30HV0) and centrifuged at 14,500 RPM for 4 minutes. The filtered homogenate was stored at -80°C until injection.

### MAT Homogenate Injection and Remating Assay

To assess the effect of MAT injection on female receptivity, 25-32 *Aedes aegypti* females per strain received a 250 nL thoracic injection of either 1x PBS (vehicle control) or MAT homogenate prepared from *Aedes aegypti* or *Aedes albopictus* males. MAT homogenate was tested at three concentrations: 1x (undiluted), 0.5x (1:1 homogenate:PBS), and 0.25x (1:3 homogenate:PBS). All injections were performed using a Nanoject III microinjector (Drummond Scientific Company) fitted with a glass needle, under a dissecting stereomicroscope. Following injection, females were grouped by treatment and transferred to small cages (21.6 cm diameter x 18.1 cm height) containing a moistened paper towel for hydration, and allowed to recover overnight at 25-28°C and 70-80% RH. After recovery, 20-30 injected females were co-housed with conspecific males at a 1:1 ratio in large rearing cages (BugDorm) under standard rearing conditions (28°C, 70-80% RH, 12 h light:12 h dark). Adults were provided 10% sucrose solution *ad libitum*. After 72 hours of co-housing, females were cold anesthetized at 4°C, and spermathecae were dissected and examined to determine insemination status.

### Statistical Analysis

All statistical analyses were performed with GraphPad Prism version 10.5 (La Jolla, CA, USA). Data are shown as medians with 95% confidence intervals unless noted otherwise. Statistical methods and sample sizes are included in figure captions. Nonparametric tests were used for data that does not follow normal distribution as determined by Shapiro-Wilk test. Statistical tests for each experiment are reported in the Figure legends and additional details are reported in **Data File S1**.

## RESULTS

### Field collection of naive and co-occurring *Aedes aegypti* and generation of lab strains

To investigate potential premating and postmating mechanisms of satyrization resistance, we established two field-derived strains of *Aedes aegypti* from Lee County, Florida: one from Fort Myers and one from Upper Captiva Island, both in the Lee County Mosquito Control District (**Figure 1A**). *Aedes aegypti* occurs throughout the region and, although *Aedes albopictus* was established in Fort Myers by 1992 [26], this species has not been detected on Gulf Coast barrier islands. Consistent with previous reports, only *Aedes aegypti* were present at collection sites on Upper Captiva Island, whereas all sites in Fort Myers included both species (**Table 1**). Both strains were maintained under laboratory conditions for multiple generations without detectable reductions in fecundity or survivorship.

### *Aedes aegypti* females from naive strain are more susceptible to insemination by *Aedes albopictus* males

Because *Aedes albopictus* co-occurs with *Aedes aegypti* at the Fort Myers collection site, females from the Fort Myers (FtM) strain are likely subject to ongoing selective pressure to evolve resistance to satyrization. In contrast, *Aedes albopictus* has not established on Upper Captiva Island, and females from the Upper Captiva (UpC) strain are therefore expected to be naïve to interspecific mating attempts and to lack satyrization resistance. To test for differences in susceptibility to satyrization between strains, 25 females from each field-derived strain were co-housed for one week with either conspecific *Aedes aegypti* males or with heterospecific *Aedes albopictus* males from the Foshan laboratory strain, after which females were scored for insemination status. Because interspecific mating is non-preferred, we tested *Aedes albopictus* males at both 1:1 and 1:2 (female:male) ratios to increase mating pressure and assess the capacity of females to resist insemination by heterospecific males under elevated exposure.

Conspecific males successfully inseminated 100% of females in both strains across all trials at the 1:1 ratio (**Figures 1B and 1C**). Interspecific mating rates were consistently lower than intraspecific rates across all conditions. UpC females showed intermediate rates of insemination by *Aedes albopictus* males at the 1:1 ratio (median = 10%), and insemination rate increased when male density was doubled (median = 24%) (**Figure 1B**). In contrast, FtM females were rarely inseminated by *Aedes albopictus* males at either 1:1 or 1:2 ratios (medians = 4% and 6% respectively) (**Figure 1C**). UpC insemination rates increased with *Aedes albopictus* male density, whereas FtM insemination rates remained consistently low regardless of male density, indicating that FtM females are more resistant to interspecific insemination than UpC females under elevated mating pressure.

### *Aedes albopictus* males make fewer successful physical contacts with co-occurring *Aedes aegypti* females

To investigate premating barriers as a potential mechanism of satyrization resistance, we used a fluorescent dye transfer assay [25] to quantify attempted mating interactions between males and females. A single fluorescent dye-painted male was co-housed with ten unpainted females, and dye transfer to the terminal abdominal segments of females was used as a proxy for physical mating interactions such as attempted copulation (**Figure 2A**). *Aedes aegypti* females from each strain were paired with either conspecific males from their own strain or heterospecific *Aedes albopictus* males. If premating barriers contribute to satyrization resistance, we predicted that FtM females would have fewer physical interactions with *Aedes albopictus* males than with conspecific males, and would show lower interspecific interaction rates than UpC females.

**Figure 2:**
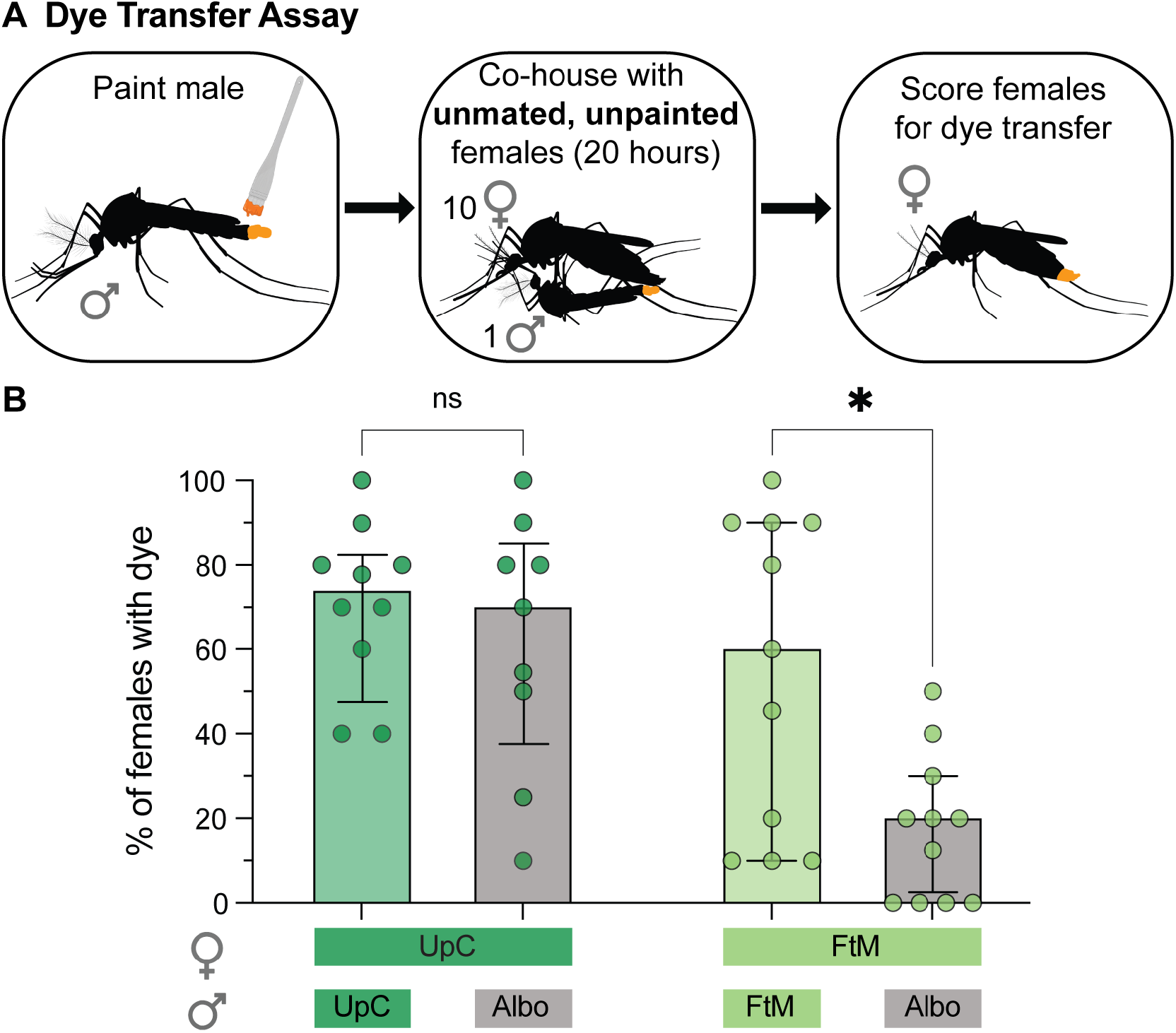
FtM females have fewer physical interactions with heterospecific males than with conspecific males, while UpC females show no difference. (A) Experimental timeline for the fluorescent dye transfer assay, in which a single dye-painted male was co-housed with 10 unpainted females. (B) Percentage of females with dye transfer to the terminal abdominal segments, used as a proxy for physical mating interactions. For all panels, n = 9-11 trials of 8-10 females per trial. Asterisks indicate statistically significant differences (p < 0.05, Mann-Whitney U test; ns, not significant). Additional statistical details provided in **Data File S1**.

Although intraspecific dye transfer rates were high in both strains, the two populations differed in their responses to heterospecific males. UpC females showed no significant difference in physical interaction rates with conspecific versus heterospecific males, whereas FtM females had significantly fewer interactions with *Aedes albopictus* males than with conspecific males (**Figure 2B**). Dye transfer rates were also significantly lower in FtM females housed with *Aedes albopictus* males than in UpC females under the same conditions (p = 0.0013, Mann-Whitney; **Data File S1**), indicating that fewer physical contacts occur between *Aedes albopictus* males and FtM females than between *Aedes albopictus* males and UpC females. Whether this reflects active rejection by females, reduced male attempts, or both remains to be determined. Insemination rates were low following intraspecific pairings in both strains, and no insemination by *Aedes albopictus* males was observed within the 20-hour assay window (**Data File S1**), likely reflecting the effects of dye application, the use of a single male, and the short duration of the assay. These results are consistent with a premating component to satyrization resistance in FtM females, reflected in reduced physical interactions with heterospecific males compared to conspecific males.

### Aedes aegypti females from co-occurring strains are less sensitive to male-derived signals from Aedes albopictus

To assess postmating barriers to remating, we tested the sensitivity of *Aedes aegypti* females to MAT homogenate injected from either conspecific *Aedes aegypti* or heterospecific *Aedes albopictus* males (**Figure 3A**). Females were injected with doses ranging from approximately one-quarter (1x) to one-sixteenth (0.25x) of a MAT. Females from both the FtM and UpC strains were highly sensitive to conspecific MAT homogenate at all three concentrations tested, showing near-complete suppression of remating even at the lowest dose (**Figure 3B**). Both strains were less sensitive to *Aedes albopictus* MAT homogenate than to conspecific MAT homogenate. However, the two strains differed in their responses to heterospecific MAT homogenate; UpC females showed greater remating suppression across all doses of *Aedes albopictus* MAT homogenate compared to FtM females (**Figure 3B**). Importantly, this reduced sensitivity in FtM females was specific to heterospecific MAT homogenate, since both strains retained strong sensitivity to conspecific male-derived signals. Together, these results indicate that FtM females differ from UpC females in their postmating physiological response to *Aedes albopictus* MAT homogenate, and reduced sensitivity to heterospecific male signals may constitute a postmating component of satyrization resistance in this population.

**Figure 3:**
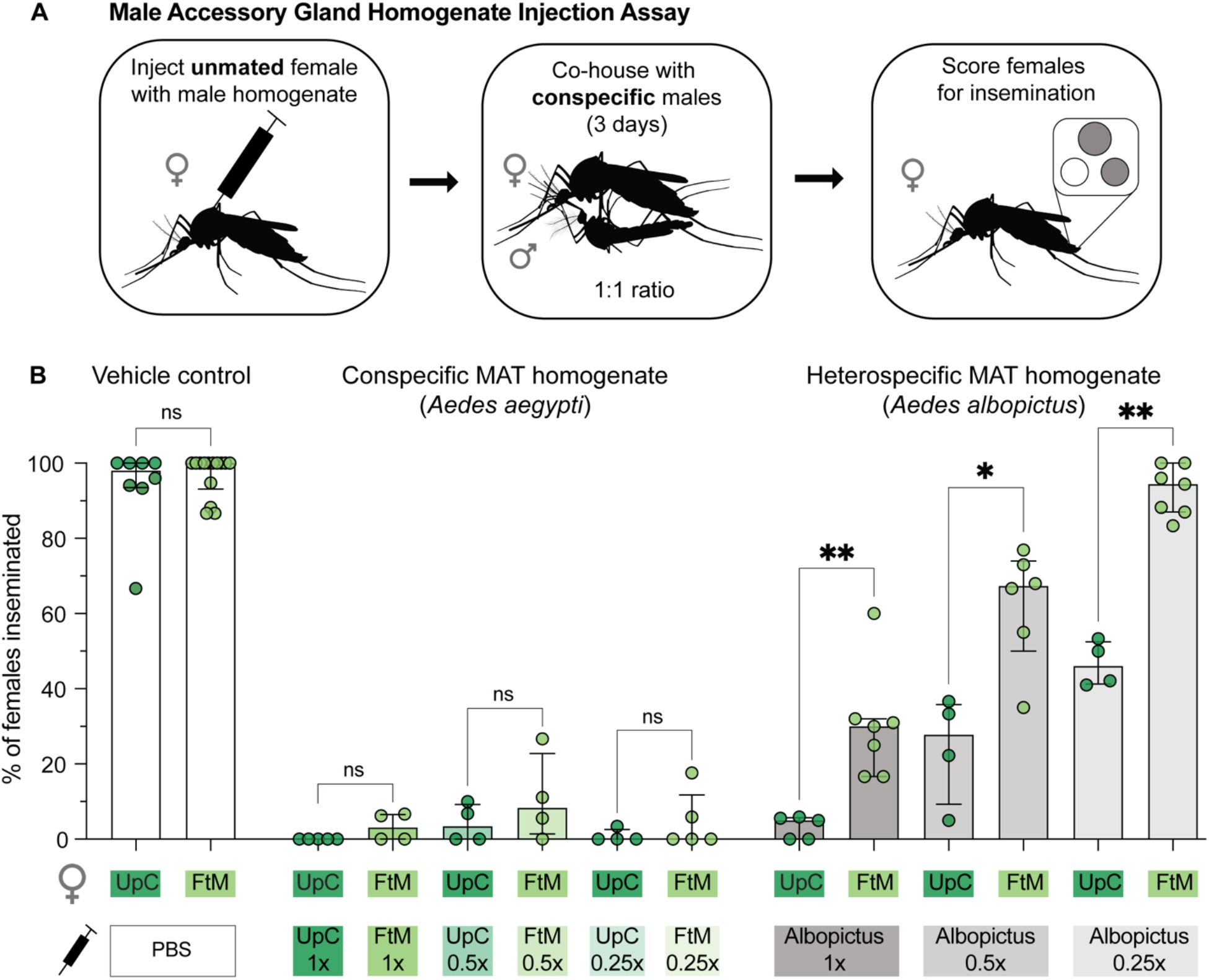
Females from co-occurring population show higher rates of remating after injection with heterospecific male abdominal tip homogenate. (A) Experimental timeline for the MAT homogenate injection and remating assay. (B) Percentage of females inseminated by conspecific males following 72 hours of co-housing after injection with MAT homogenate from the indicated male source at the indicated concentration. n = 4-14 trials with 21-31 females per trial. (* = p < 0.05; ** = p < 0.01; Mann-Whitney U test with Benjamini-Hochberg false discovery rate correction; ns, not significant). Additional statistical details provided in **Data File S1**.

## DISCUSSION

In this study, we identified premating and postmating differences between *Aedes aegypti* strains associated with co-occurrence with *Aedes albopictus*. Females from the FtM population, which co-occurs with *Aedes albopictus* in the field, showed fewer physical interactions with heterospecific males, lower rates of interspecific insemination, and higher rates of intraspecific remating following injection with heterospecific MAT homogenate, compared to females from the naive UpC population. It is important to note that the experimental conditions used here, including elevated male densities and direct MAT homogenate injection, do not reflect the natural dynamics of field encounters between these species. Instead, these assays were designed as laboratory “stress tests” intended to maximize mating pressure and isolate specific components of the mating interaction, to reveal biological differences between strains that might not be detectable under more naturalistic conditions. This approach allows us to generate mechanistic hypotheses about the basis of satyrization resistance that can be tested in future studies. Within this framework, our results suggest that FtM females are better able to avoid interspecific mating attempts, and that paternity-enforcing signals derived from *Aedes albopictus* males are less effective at suppressing remating in females from the co-occurring population. Even under elevated male densities, FtM females remained largely resistant to insemination by *Aedes albopictus* males. Similarly, when injected with heterospecific MAT homogenate, FtM females remated with conspecific males at higher rates than UpC females, and this difference became more pronounced at lower homogenate concentrations. In contrast, UpC females were more susceptible to heterospecific insemination during co-housing and showed greater suppression of remating following heterospecific MAT injection, consistent with a lack of resistance in this population.

Together, these findings implicate both enhanced mate discrimination and reduced sensitivity to heterospecific male signals as components of satyrization resistance in FtM females. Our results are consistent with previous work demonstrating that multigenerational exposure to *Aedes albopictus* can favor the development of resistance behaviors in *Aedes aegypti* [14] and extend those findings by identifying distinct premating and postmating mechanisms that may contribute to this resistance.

### Potential mechanisms of premating barriers

The mechanisms underlying mate recognition in *Aedes aegypti* remain poorly understood despite their critical importance for female fitness. During mating interactions, multiple sensory modalities are likely involved, including visual signals, wingbeat frequency, and chemical communication. Most mating attempts are unsuccessful, and females employ a range of active rejection behaviors, including evasive flight, kicking, abdominal twisting, and failure to open the vaginal plates, to avoid insemination [15,27–29]. FtM females may adjust their mating receptivity to more effectively reject *Aedes albopictus* males while maintaining normal receptivity toward conspecific males, suggesting that co-occurrence with *Aedes albopictus* has shaped female discrimination abilities in this population.

Female wingbeat frequency is widely regarded as a primary sensory cue driving male attraction and mate selection [15,27,30,31]. Females of the two species differ acoustically, with *Aedes aegypti* females producing louder sounds with more harmonics than *Aedes albopictus* females [32]. Additionally, studies suggest that there are differences between these species in their reliance on acoustic cues: *Aedes aegypti* males respond strongly to female flight sounds by orienting and attempting copulation, whereas *Aedes albopictus* males show weaker responses to flight sounds from either species [33]. These acoustic differences may contribute to premating discrimination, though the extent to which females also use acoustic cues to assess male identity remains unclear.

Chemical signals represent another plausible mechanism through which females could discriminate between conspecific and heterospecific males prior to mating. Several male- and female-derived pheromones have been proposed to promote aggregation in *Aedes aegypti* [34], support mating success in *Anopheles stephensi* [35], or contribute to reproductive isolation in *Aedes albopictus* [36]. Species-specific pheromones could play a role in the development of satyrization resistance, as a similar effect has been observed in *Drosopholids* [37]. In *Drosophila melanogaster* and *Drosophila simulans*, two closely related species of fruit fly that frequently co-occur in the wild, interspecific mate avoidance is regulated by the recognition of species-specific pheromones produced by females, which selectively promote male courtship towards conspecific females [38]. An analogous mechanism in *Aedes aegypti* would be consistent with our observation that FtM females show reduced physical interactions specifically with heterospecific males. Additionally, premating barriers are unlikely to operate exclusively through female behavior; recent work has demonstrated that males also terminate mating interactions, and this is particularly common in interspecific encounters [39], suggesting that the reduced physical contact reflects contributions from both sexes.

### Potential mechanisms of postmating differences

FtM females showed a striking pattern of selectivity in their responses to MAT homogenate; they remated with conspecific males at high rates following injection with *Aedes albopictus* MAT homogenate, yet remained strongly suppressed from remating after injection with conspecific *Aedes aegypti* MAT homogenate. This indicates that FtM females are capable of differentially responding to paternity-enforcing signals from the two species, effectively ignoring heterospecific signals while remaining sensitive to conspecific ones. This selectivity implies that resistance does not reflect a generalized reduction in sensitivity to male-derived signals, but rather a specific change in how FtM females detect or respond to *Aedes albopictus*-derived compounds, raising the question of which molecular components of MAT homogenate are responsible and the possibility that their cognate receptors differ in specificity between the two populations.

The relevant signals are contained within male seminal fluid proteins (SFPs), which modulate postmating female physiology and behavior across a wide range of insect species [40,41]. One example in *Aedes aegypti* is Head Peptide-I (HP-I), a male-derived peptide that can activate Neuropeptide Y-Like Receptor 1 (NPYLR1) in females to enforce male paternity within a few hours of mating [42,43]. Notably, *Aedes albopictus* HP-I peptides also activate *Aedes aegypti* NPYLR1 [43], confirming that heterospecific signals are capable of activating the same receptor in *Aedes aegypti*. However, because HP-I acts only transiently, the long-term paternity enforcement observed following MAT injection involves additional, as yet unidentified ligands and receptors. The identities of male-derived signals responsible for lifelong paternity enforcement and their corresponding female receptors remain unknown. Thousands of proteins have been identified in the *Aedes aegypti* male ejaculate, and SFP-associated proteins are more highly enriched for signal peptides than those associated with sperm or testis [44,45] making seminal fluid composition a promising target for future investigation. Studies across insect species underscore both the importance and the diversity of SFP-mediated postmating responses in insects. The *Drosophila* Sex Peptide remains the best-characterized example, and active efforts to characterize seminal fluid composition across species, including recent work in brown planthoppers and moths [46–49] highlight the generality of this mechanism. Notably, paternity enforcement in *Anopheles* mosquitoes is mediated by a steroid hormone rather than a protein, [50] illustrating that even within dipterans, the molecular substrates of postmating responses are diverse. These comparisons suggest that the receptors and signaling pathways mediating paternity enforcement in *Aedes aegypti* females may be more complex than currently appreciated, and that population-level variation in these pathways presents a tractable entry point for future mechanistic studies.

### Implications for vector control

Our findings have direct relevance for vector control strategies that depend on mating interactions. Sterile male release programs and related approaches rely on the assumption that target females will mate with released males and subsequently fail to reproduce. However, if wild *Aedes aegypti* populations in areas of *Aedes albopictus* co-occurrence have gained both behavioral and physiological resistance to interspecific mating attempts that alter their interactions with conspecific males, this could reduce the efficacy of control programs. More broadly, our results highlight that mating behavior in *Aedes aegypti* is not fixed but can vary meaningfully between populations in ways that reflect local evolutionary history. Understanding this natural variation, its mechanistic basis, heritability, and distribution across wild populations will be essential for the design and evaluation of mating-based vector control strategies. The premating and postmating mechanisms identified here provide concrete targets for future work aimed at characterizing the molecular and sensory basis of satyrization resistance in this important disease vector.

## Supporting information

DataS1

## Acknowledgements

We thank members of the Duvall Lab for their critical comments on the manuscript. We thank Rachel Morreale, Stephen Stenhouse, Dr. David Hoel, and Aaron Lloyd at Lee County Mosquito Control District for their guidance in selecting trapping sites and contributions to our field collecting efforts. We thank Monica Cramer for her contribution to field collecting. This work was supported by the following grants: NIGMS (R35 GM137888) (LBD), Beckman Young Investigator Award (LBD), Pew Scholar in Biomedical Sciences Award (LBD).

## Supporting Information

**Data File S1**. This file includes all raw data included in the manuscript and additional statistical details.

## Author Contributions

**Conceptualization:** Lauren E. Subramaniam, Thomas M. Gabel, Laura B. Duvall

**Data curation:** Lauren E. Subramaniam, Olivia M. Martinez, Thomas M. Gabel, Laura B. Duvall

**Formal Analysis:** Lauren E. Subramaniam, Olivia M. Martinez, Thomas M. Gabel, Laura B. Duvall

**Funding Acquisition:** Laura B. Duvall

**Investigation:** Lauren E. Subramaniam, Olivia M. Martinez, Thomas M. Gabel

**Methodology:** Lauren E. Subramaniam, Olivia M. Martinez, Thomas M. Gabel

**Project Administration:** Thomas M. Gabel, Laura B. Duvall

**Resources:** Thomas M. Gabel

**Supervision:** Laura B. Duvall

**Validation:** Olivia M. Martinez, Thomas M. Gabel

**Visualization:** Lauren E. Subramaniam, Olivia M. Martinez, Thomas M. Gabel, Laura B. Duvall

**Writing – original draft:** Lauren E. Subramaniam, Thomas M. Gabel, Laura B. Duvall

**Writing – review & editing**: Lauren E. Subramaniam, Olivia M. Martinez, Thomas M. Gabel, Laura B. Duvall

